# Cochlear theta activity oscillates in phase opposition during interaural attention

**DOI:** 10.1101/2022.02.21.481289

**Authors:** Moritz Herbert Albrecht Köhler, Nathan Weisz

**Author notes:** **Corresponding author**, (MHA.K.).

## Abstract

It is widely established that sensory perception is a rhythmic process as opposed to a continuous one. In the context of auditory perception this effect is only established on a cortical and behavioral level. Yet, the unique architecture of the auditory sensory system allows its primary sensory cortex to modulate the processes of its sensory receptors at the cochlear level. Previously, we could demonstrate the existence of a genuine cochlear theta (~6 Hz) rhythm that is modulated in amplitude by intermodal selective attention. As the study’s paradigm was not suited to assess attentional effects on the oscillatory phase of cochlear activity the question whether attention can also affect the temporal organization of the cochlea’s ongoing activity remained open. The present study utilizes an interaural attention paradigm to investigate ongoing otoacoustic activity during a stimulus-free cue-target interval and an omission period of the auditory target in humans. We were able to replicate the existence of the cochlear theta rhythm. Importantly, we found significant phase opposition between the two ears and attention conditions of anticipatory as well as cochlear oscillatory activity during target presentation. Yet, the amplitude was unaffected by interaural attention. These results are the first to demonstrate that intermodal and interaural attention deploy different aspects of excitation and inhibition at the first level of auditory processing. While intermodal attention modulates the level of cochlear activity, interaural attention modulates the timing.

## Introduction

A large body of literature states that sensory perception is a rhythmic process rather than a continuous one. This is largely based on studies of brain rhythms that reflect the synchronized modulations of excitability of large ensembles of neurons (Kayser et al., 2015; Lakatos et al., 2005; Romei et al., 2008). Further evidence can be found on a behavioral level, in domains such as object exploration (Wöstmann et al., 2016; Wyart et al., 2012) or attentional modulations of perception (Busch & VanRullen, 2010; Fiebelkorn et al., 2013). Frequencies in the range of the cortical theta-band (~3-8 Hz) have also come into focus as being temporal organizers of perception.

Influential frameworks established on research in the visual modality suggest that an attention network operating in the theta frequency range assists in the temporal organization of neural activity associated with perception and action (Fiebelkorn & Kastner, 2019; Landau & Fries, 2012). Hence, large-scale theta rhythms possibly aid in preventing conflicts between sensory and motor functions. Beside modulations in power and frequency, according to these frameworks, the phase of theta rhythms putatively plays an essential role for how attentional sampling is deployed. For example, Fiebelkorn & Kastner (2019) proposed the existence of two alternating states that coordinate sensory and motor processes via phase opposition. That is, the phase of the theta rhythm encourages either sampling of a relevant feature (such as location) or shifting to another.

On a general level, attentional rhythms should also exist in other sensory modalities (e.g., the auditory modality) to similarly help in the coordination of perception and action. Mounting evidence suggests a key role of theta also for auditory perception at various levels of complexity (e.g. for simple target detection (Ng et al., 2012); speech perception (Poeppel & Assaneo, 2020); auditory scene exploration (Kayser, 2019)). Albeit early studies failed to provide evidence for the existence of thetarhythmic oscillations in auditory attention (İlhan & VanRullen, 2012; Zoefel & Heil, 2013), recent findings from both cortical and behavioral data speak for their existence (Ho et al., 2017, 2019; Kayser, 2019; Kubetschek & Kayser, 2021; Ng et al., 2012; Plöchl et al., 2021). Interestingly, in bilateral pitch-identification tasks behavioral performance theta-rhythmically oscillates in antiphase between the two ears (Ho et al., 2017; Plöchl et al., 2021). These findings hint to the existence of a similar theta-dependent mechanism in auditory attention as proposed for visual attention.

The interpretations of the evidence for rhythmic perception in the auditory modality are largely cortico-centric. Yet, the auditory system is unique with respect to other modalities in that its neuronal architecture allows its primary sensory cortex to modulate the activity of its sensory receptors at the cochlear level. Indeed, mainly by measuring otoacoustic emissions (Dragicevic et al., 2019; Giard et al., 1994; Marcenaro et al., 2021; Wittekindt et al., 2014), it has been established that attention affects cochlear properties. Other approaches such as cochlear microphonics (Delano et al., 2007) or direct measurement of hearing nerve activity (Gehmacher et al., 2022) support this notion, which can normally only be collected in very rare circumstances in humans (e.g., individuals with cochlear implants). Despite its easy applicability, otoacoustic emissions (OAE) are restricted to sound-evoked responses, which are likely affected by confounding medial olivocochlear (MOC) activity (Guinan et al., 2003). Thus, they only provide an extremely limited view of (top-down) oscillatory dynamics at the cochlear level.

In a previous study we introduced the measurement of ongoing otoacoustic activity (OOA) during silent cue-target intervals (Köhler et al., 2021). The measurement of OOA offers various advantages over OAEs that include the prevention of unwanted MOC activity, the possibility to analyze ongoing temporal patterns of cochlear acoustic activity, and the implementation of attention paradigms that stay very close to the cortical literature. From this, we could demonstrate the existence of a genuine cochlear theta (~6 Hz) rhythm that is modulated in amplitude but not frequency nor phase by intermodal selective attention (Köhler et al., 2021). However, the study’s paradigm was not suited to assess possible attentional effects on the oscillatory phase of OOA.

The current study aims to shed more light on the properties of the oscillatory activity at the most peripheral stage of the auditory system. Thus far, we could demonstrate that attention modulates the overall level of cochlear activity (Köhler et al., 2021). Whether and how cochlear activity is temporally orchestrated (e.g. expressed by oscillatory phase modulations) still remains unknown. Staying close to our previous study, we implemented an interaural attention paradigm,which aimed to assess attentional phase effects on the cochlear level.

Replicating the findings from our previous study we found a dominant theta rhythmicity of cochlear activity. By contrast to intermodal attention (Köhler et al., 2021), interaural attention modulates the phase of OOA. Furthermore, these signals are not linked to eye movement-related putative eardrum oscillations (Gruters et al., 2018). Therefore, our results demonstrate that attentional processes can impact the timing of cochlear activity. Interestingly, this phase opposition is present during anticipation of a target as well as its presentation.

## Materials & Methods

### Participants

31 healthy volunteers (21 females, age range: 18-41 years) participated in this study. After finishing the experiment, one participant reported that she was not motivated and tried to finish the experiment as fast as possible. As a result, this participant was excluded from analyses. None of the subjects reported any known hearing deficit and any visual impairment was sufficiently corrected. Moreover, all subjects were instructed to not take any ototoxic drugs and expose themselves to any loud noise 24 hours before their participation in the experiment. Subjects were informed about the experimental procedure and the purpose of the study and gave written informed consent. As compensation, subjects received either 10 euro per hour or credit for their psychology studies. This study was approved by the Ethics Committee at the University of Salzburg.

### Stimuli and Procedure

The study’s focus was to investigate interaural attention processes at the level of the cochlea during a silent cue-target interval. Previously, numerous studies used a block design to investigate attentional modulations of OAEs and were criticized for not achieving highly controlled attentional conditions (Carrasco et al., 2004; Köhler et al., 2021; Ward, 1997; Wittekindt et al., 2014). Therefore, we implemented a trial-wise cueing paradigm.

Measurements took place in an EEG-recording room, in which subjects sat alone and quietly in front of a PC screen (screen ratio: 16:9; width: ca. 56 degrees of visual angle; refresh rate: 120 Hz). Participants performed five blocks consisting of 100 trials each and six experimental conditions in total. The experimental conditions were defined by (1) the presented cue and (2) the congruency of the target. In the present study only two conditions are of relevance. **Figure 1** schematically illustrates the course of each trial. Each trial started with a visually presented cue (1 s duration) instructing the participants to shift their attention to their left and right ear (below referred to as Attend Left & Attend Right conditions), respectively. The direction, to which the presented arrow pointed, specified the to-be-attended ear. After cue presentation a fixation cross was shown for 2 s. During this silent cue-target interval subjects had to actively shift their attention to the indicated ear. By design the cue was 100% informative. This ensured that any effects of divided attention were avoided and subjects’ focus was completely placed on the cued ear (Köhler et al., 2021; Wittekindt et al., 2014). The target stimulus was in all conditions a bilaterally presented click-train (duration: 500 ms; click-frequency: 24 Hz; loudness: 64 dB SPL) consisting of 14 clicks of 80 μs each. In 80 trials of each block there was an omission of three clicks in either the left or right ear. The occurrence of the omissions was counterbalanced across both ears resulting in 40 trials with an omission in the left ear and 40 trials in the right ear. Moreover, the beginning of the omissions was pseudorandomly jittered between the fourth and tenth click. The task was to identify if there was an omission in the to-be-attended ear. Next, a response screen, indicated by a centrally placed question mark, was shown for 2 s. Participants were instructed to respond as fast and correctly as possible by pressing the corresponding key of the keyboard (“N” & “M”) with their index and middle finger of one hand, respectively. The used hand was counterbalanced across all subjects. The inter-trial interval was jittered uniformly between 1 and 2 s. Experimental stimulation was implemented by use of the Psychophysics Toolbox Version 3 (Brainard, 1997) and the Objective Psychophysics Toolbox (Hartmann & Weisz, 2020), which provides an intuitive, unified, and clear interface on top of the Psychophysics Toolbox, using custom-written MATLAB scripts (Version 9.8; The MathWorks, Natick, Massachusetts, USA).

**Figure 1.**
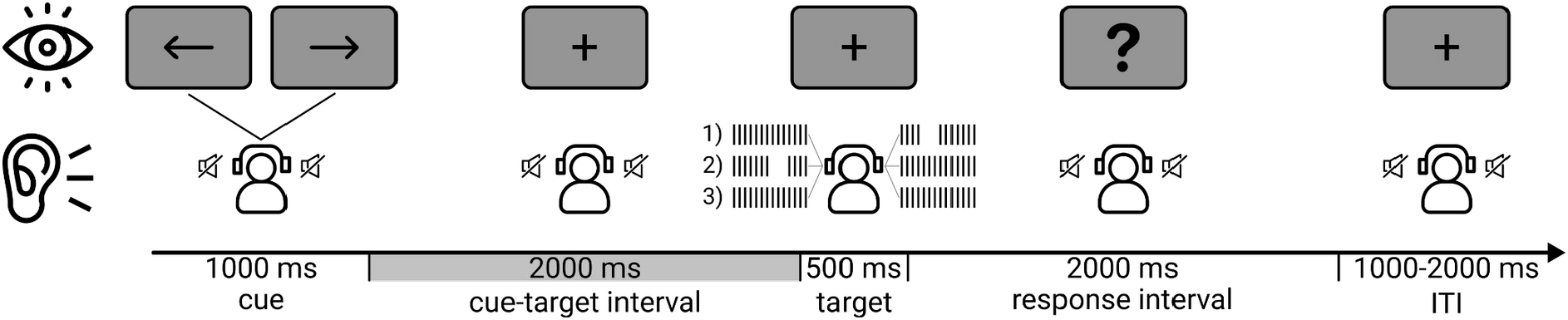
Schematic illustration of the task. Each trial started with a cue (100% informative) instructing participants to shift their attention to their left or right ear. A silent cue-target interval, in which participants focused their attention, followed. Thereafter the target was presented. The target consisted of a click train (24 Hz presentation rate) that was presented bilaterally while in 80% of the trials there was an omission of three clicks in one ear. The onset of the omission periods was jittered between the 3^rd^ and 10^th^ click of 14 clicks in total. Subsequently, participants indicated by a button press on the keyboard if the omission period occurred in the to-be-attended ear. The inter-trial interval was jittered uniformly between 1 and 2 s.

### Apparatus

#### Ongoing Otoacoustic Activity

For every subject OOA was measured by a probe consisting of a sensitive microphone and two loudspeakers (ER-10C microphone/preamplifier system, Etymotic Research, Elk Grove Village, Illinois, USA). After cleaning each ear canal from excessive cerumen a probe was fitted with a foam ear tip that sealed the ear canal. OOA was recorded from both ears concurrently without preamplification. The microphone signal was fed into a g.USBamp Research (g.tec medical engineering GmbH, Schiedlberg, Austria) with a sampling rate of 38.4 kHz. The two ER-10C received their input via a 3.5 mm audio jack to BNC cable coming from a sound preamplifier (SLA-35 stereo matching amplifier, Monacor International, Bremen, Germany), which again received its input from a computer sound card (Realtek Semiconductors Corp., Hsinchu, Taiwan). Event triggering was performed by a U3-LV (LabJack Corporation, Lakewood, Colorado, USA).

#### Electroencephalography

Electroencephalography (EEG) was recorded for a separate study question. However, the data was used to address a concern linked to the relation of our OOA results to eye movements. For this purpose 28 active electrodes (g.LADYbird, g.tec medical engineering GmbH, Schiedlberg, Austria) were attached to a g.GAMMAcap (g.tec medical engineering GmbH, Schiedlberg, Austria). The electrodes were evenly distributed across the cap and placed on the following positions of the international 10-10 system: Fpz, F7, F3, Fz, F4, F8, FC5, FC1, FC2, FC6, C3, Cz, C4, T7, T8, CP5, CP1, CP2, CP6, P7, P3, Pz, P4, P8, POz, O1, Oz, and O2. The reference electrode was placed on the right ear lobe and AFz was used as a ground. Impedance was kept below 5 kΩ in all subjects for the whole experiment. The signals were fed into two g.USBamp Research (g.tec medical engineering GmbH, Schiedlberg, Austria) via two g.GAMMAboxes (g.tec medical engineering GmbH, Schiedlberg, Austria) with a sampling rate of 38.4 kHz.

### Signal Processing

#### Ongoing Otoacoustic Activity

We preprocessed the raw data with the open-source FieldTrip toolbox for M/EEG data (Oostenveld et al., 2011) and custom-written MATLAB scripts (Version 9.8; The MathWorks, Natick, Massachusetts, USA).

First, we high-pass filtered the raw signal at 500 Hz (kaiser-windowed sinc FIR filter, onepass-zerophase, order: 1114, transition width: 125 Hz) to eliminate any influence of “low-frequency” noise on otoacoustic activity. Moreover, otoacoustic activity is routinely found in a frequency range of 500 - 4000 Hz (Froehlich et al., 1993; Meric & Collet, 1992, 1994).

For the analysis in the cue-target interval we split the filtered data into 2 s trials that contained the signal from the cue-target intervals. We identified bad trials that contained “high-frequency” noise (e.g. caused by moving, swallowing, and coughing) by Hilbert-transforming and z-scoring the signal. We excluded a trial from further analyses of the cue-target interval if the proportion of samples that exceeded a z-score of 8 was above 1%. On average we excluded 8.16% of the trials (*SD* = 5.97%) per subject.

For the analysis in the omission period we split the filtered data into 125 ms trials with the signal from the omission period. We shortened the originally calculated trial length of 162.4 ms (4 x 40 ms gaps between clicks + 3 x 80 μs clicks) by 37.4 ms to compensate for trigger inaccuracies and ringing of the click tones. There was no trial rejection as the trials were very short and noise was not problematic in the omission periods.

In a last step we separately extracted for the 2 s and 125 ms trials the absolute part of the Hilbert transform by applying a bandpass filter between 1500 and 2000 Hz (kaiser-windowed sinc FIR filter, onepass-zerophase, order: 372, transition width: 375 Hz) as otoacoustic activity is particularly pronounced in this frequency range (Puria, 2003). By means of the absolute part of the Hilbert transform the envelope of the OOA is extracted. The envelope represents amplitude fluctuations of outer hair cell (OHC) activity at frequencies of the bandpass window (Köhler et al., 2021). While raw OHC activity is found at frequencies (500 - 4000 Hz) that are significantly beyond that of cortical oscillatory activity, which is pronounced between 1 and 80 Hz, the extraction of the envelope allows analyses of cochlear modulations at frequencies that are commonly used in evaluations of cortical activity during cognitive tasks. This approach facilitates the integration of OHC modulatory activity at specific acoustic frequencies and cortical activity. All further analyses are based on the envelope of the OOA.

#### Electroencephalography

We conducted preprocessing of the EEG data with the help of the NoiseTools toolbox (de Cheveigné & Arzounian, 2018), the open-source FieldTrip toolbox for M/EEG data (Oostenveld et al., 2011) and custom-written MATLAB scripts (Version 9.8; The MathWorks, Natick, Massachusetts, USA). At first, the raw data was downsampled to 3.84 kHz and channels that comprised more than 40% samples that exceeded three times the median absolute value over all data were marked as bad and excluded from further analysis. We detrended the data by removing the 10th-degree polynomial trend. In order to boost the reliability of the later calculated independent component analysis (ICA) we applied a 40 Hz low-pass filter (kaiser-windowed sinc FIR filter, onepass-zerophase, order: 1392, transition width: 10 Hz) and a 0.1 Hz high-pass filter (kaiser-windowed sinc FIR filter, onepass-zerophase, order: 69542, transition width: 0.2 Hz). We split the EEG data into the same trials as the OOA data and only kept trials that were marked as good for the OOA data. Next, we applied an average rereference. Lastly, we conducted an ICA and identified components that in terms of topography and time course clearly depict vertical and horizontal eye movements. These components were used to link ocular activity to OOA (see below).

### Induced Power Analysis

We conducted induced power analysis only for the data from the cue-target interval. To calculate induced power spectral density (PSD) we zero-padded the envelope signal from the 2 s cue-target interval to a length of 3.4133 s. Next, we calculated a fast fourier transform with multiple tapers (“mtmfft” FieldTrip implementation) from discrete prolate spheroidal sequences (dpss) including a frequency range of 1 - 45 Hz with a bin size of 0.5 Hz and a frequency smoothing of 1 Hz. As a result we obtained four PSDs - from both ears and the two experimental conditions (Attend Left & Attend Right) - per subject.

In a next analysis step we parameterized the individual PSDs of the OOA envelope by the usage of the FOOOF-toolbox (Donoghue et al., 2020) in Python (Version 3.8.1). The advantage of FOOOF is that it characterizes putative oscillations in neural power spectra without aperiodic contributions. The frequency range of the parameterization was set from 1 - 20 Hz. We left all parameters at their respective defaults except the value of the peak threshold, which we set to one standard deviation (SD).

### Evoked Power Analysis

Besides induced power we also calculated evoked power from the omission periods. By doing so we were able to analyze the magnitude of the contralaterally evoked OAEs during stimulus presentation. We did not apply zero-padding as the frequency resolution of 8 Hz for the 125 ms interval perfectly fitted the 24 Hz stimulation rate of the click train. We calculated a fast fourier transform with multiple tapers (“mtmfft” FieldTrip implementation) from discrete prolate spheroidal sequences (dpss) including a frequency range of 0-32 Hz with a bin size of 8 Hz and a frequency smoothing of 8 Hz.

### Phase Opposition Analysis

We conducted phase analysis for both the data from the cue-target intervals and omission periods. For both signals we zero-padded the trial lengths to 3.4133 s to ensure comparability between each other by obtaining the same frequency bins. Afterward, we extracted the fourier coefficients by calculating a fast fourier transform with a Hanning taper (“mtmfft” FieldTrip implementation) from 1 - 45 Hz with a bin size of 0.5 Hz and no frequency smoothing. Analogue to the power analysis we obtained four fourier spectra per subject.

In order to assess effects of phase differences between ears and interaural attention conditions we calculated the phase opposition sum (POS), introduced by VanRullen (2016), for all possible contrasts. The POS is a measure for the consistency of phase differences over trials and is based on the magnitude of the inter-trial coherence (ITC) over all trials and each experimental condition. **Figure 4A** schematically illustrates the rationale of this type of analysis. Phase opposition is described as the difference in angles for two signals that oscillate at the same temporal frequency. At time points with a 180° phase difference it reaches its maximum. The following formula shows how the POS is calculated:

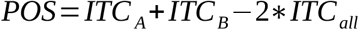

The POS will be positive when the summed ITC of each group is higher than two times the overall ITC.

### Phase Locking Analysis Between Eye Movements and OOA

In case OOA (as operationalized in this study) was driven by eye movements, then respective signals should be strongly phase coupled.

We identified for every subject the ICA-component that included activity induced by eye movements and extracted the fourier coefficients the same way as described in the section above. Again, we obtained four fourier spectra per subject. To assess phase synchrony between the signal from eye movements and the signal from the left and right ear we calculated the phase locking value (PLV) during the cue-target interval (Lachaux et al., 1999). It is defined by the following formula:

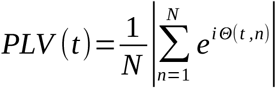

The PLV measures the inter-trial variability of the phase difference between two signals at a given frequency. It is close to 1 if the relative phase is identical across trials and 0 if there is no phase synchrony.

## Statistical Analysis

### Periodic Component Analysis

We conducted a permutation procedure to statistically assess if the periodic components that FOOOF identified differ from a null effect of their respective spectra. For this analysis we calculated 10000 permutations of the time series of the Hilbert transform and calculated the power spectra in the same way as for the real data. Finally, we compared the distribution of the surrogate power values to the real power value. We would reject the null hypothesis that the periodic component is generic when its value was equal or above the 95^th^-percentile of the surrogate distribution. Statistical analysis was performed in MATLAB (Version 9.8; The MathWorks, Natick, Massachusetts, USA) with custom-written scripts.

### Induced Power Analysis

First, we excluded all subjects where FOOOF failed to identify a peak in one of their four models. Secondly, we excluded all subjects that showed a bad model fit in one of their models. We classified a fit as bad when its value was at least two SDs smaller than the overall average across all models. Finally, we identified outliers in the distributions of each dependent variable by the use of Tukey’s boxplot rule. In total we included 19 subjects for the statistical analysis of peak frequency and 16 subjects for peak height. After these procedures, we calculated two factorial ANOVAs (2×2) with the repeated measures factors ear (left & right) and condition (Attend Left & Attend Right) for peak frequency and height that were extracted by FOOOF. All statistical analyses were performed using the R packages “ez” (Lawrence, 2016), “DescTools” (Signorell, 2021), and custom-written R scripts (Version 4.1.2; R Core Team, 2021).

### Evoked Power Analysis

For the statistical analysis of differences in evoked power between the attention conditions at the stimulation rate of the click train we excluded outliers in the distributions of the dependent variable by the use of Tukey’s boxplot rule. For the power from the left ear 27 subjects and for the right ear 25 subjects were included. Subsequently, we compared the differences by two dependent-samples t-tests, whose results were corrected for multiple comparisons using false detection rate (Benjamini & Hochberg, 1995).

### Phase Opposition & Phase Locking Analysis

We assessed statistical significance for each subject’s POS values and PLV analogously by the use of a permutation approach. We calculated 1000 surrogate POS values where for every permutation trials were pseudorandomly assigned to one of the two contrasted conditions. To calculate the 1000 surrogate PLVs the trial order of the eye movement activity was pseudorandomly permuted. In the next step we calculated the proportion of surrogate POS values / PLVs that were higher than the POS value / PLV from the original data. We set the threshold for statistical significance of the permutation p-values at 5 %. After the calculation of single subject p-values, we combined the p-values across subjects to test for the presence of a group-level effect. For this combination we used Worsely and Friston’s Method where for every frequency bin the largest p-value across subjects is elevated to the power of N (Worsley & Friston, 2000). This method is considered rather stringent as only the highest p-value across subjects influences the combined p-value. As a result a combined p-value below the 5%-significance level implies that the p-values in every single subject are at or below that level. Following this, we used an empirical adaptation of Brown’s Method to combine the p-values of frequency bins (Brown, 1975; Poole et al., 2016). This adaptation uses the empirical cumulative distribution function derived from the data instead of numerical integration (Poole et al., 2016). Statistical analysis was performed in MATLAB (Version 9.8; The MathWorks, Natick, Massachusetts, USA) with off-the-shelf functions provided by VanRullen (2016), Poole et al. (2016), and custom-written scripts.

## Results

### Behavioral Accuracy Does Not Differ Between Attention Conditions

The overall performance in terms of accuracy was *M* = 73.21% (*SD* = 22.50%). The accuracy for the Attend Left condition was *M* = 72.99% (*SD* = 22.57%) and for the Attend Right condition *M* = 73.44% (*SD* = 23.07). There was no significant difference between both conditions (*T*_(29)_ = −0.33, *p* = 0.75).

### OOA Power and Frequency at Theta Rhythm Is Not Modulated by Interaural Attention

The existence of a general endogenous cochlear rhythm in the theta range has been suggested whereby the rhythm’s frequency was not modulated by intermodal (visual vs. auditory) selective attention and did not differ between ears (Köhler et al., 2021). Moreover, the rhythm’s amplitude was increased when attention was directed to the auditory compared to the visual modality. Hence, we asked the question if theta-rhythmic activity of the cochlea is also present during interaural attention processes. Secondly, we hypothesized that the amplitude of the cochlea’s theta rhythm is enhanced in the to-be-attended ear compared to the to-be-ignored ear.

Analogous to previous literature we defined the silent cue-target interval as the period in which interaural attention processes occur (Köhler et al., 2021; Wittekindt et al., 2014). In this interval we parameterized induced cochlear oscillatory power by the usage of FOOOF (for details see the METHODS section). FOOOF could identify a peak in all but six power spectra. The average peak frequency in the left ear for the Attend Left condition was at *M* = 4.96 Hz (*SD* = 2.89 Hz) and in the Attend Right condition at *M* = 5.34 Hz (*SD* = 3.50 Hz). In the right ear the average peak frequency in the Attend Left condition was at *M* = 5.98 Hz (*SD* = 3.40 Hz) and in the Attend Right condition at *M* = 4.33 Hz (*SD* = 3.08 Hz). By means of a permutation procedure we tested if the peaks that were identified by FOOOF significantly differed from a null effect. Indeed, for all subjects every identified peak was significantly different from being a generic component. These results replicate the findings from previous studies (Dragicevic et al., 2019; Köhler et al., 2021) and contribute further evidence for the existence of a genuine cochlear theta rhythm.

Next, we tested if the frequencies in each condition and ear are significantly different. The result of a two factorial ANOVA (2×2) with the repeated measures factors ear (left & right) and attention condition (Attend Left & Attend Right) revealed no significant main effects (ear: *F*_(1, 18)_ = 5.08e-05, *p* = 0.99; condition: *F*_(1, 18)_ = 0.78, *p* = 0.39). Regarding the interaction, **Figure 2A** suggests a deceleration of the cochlear’s theta rhythm for the to-be-attended and acceleration for the to-be-ignored ear. However, the effect was only significant on a trend level (*F*_(1, 18)_ = 3.93, *p* = 0.06).

**Figure 2.**
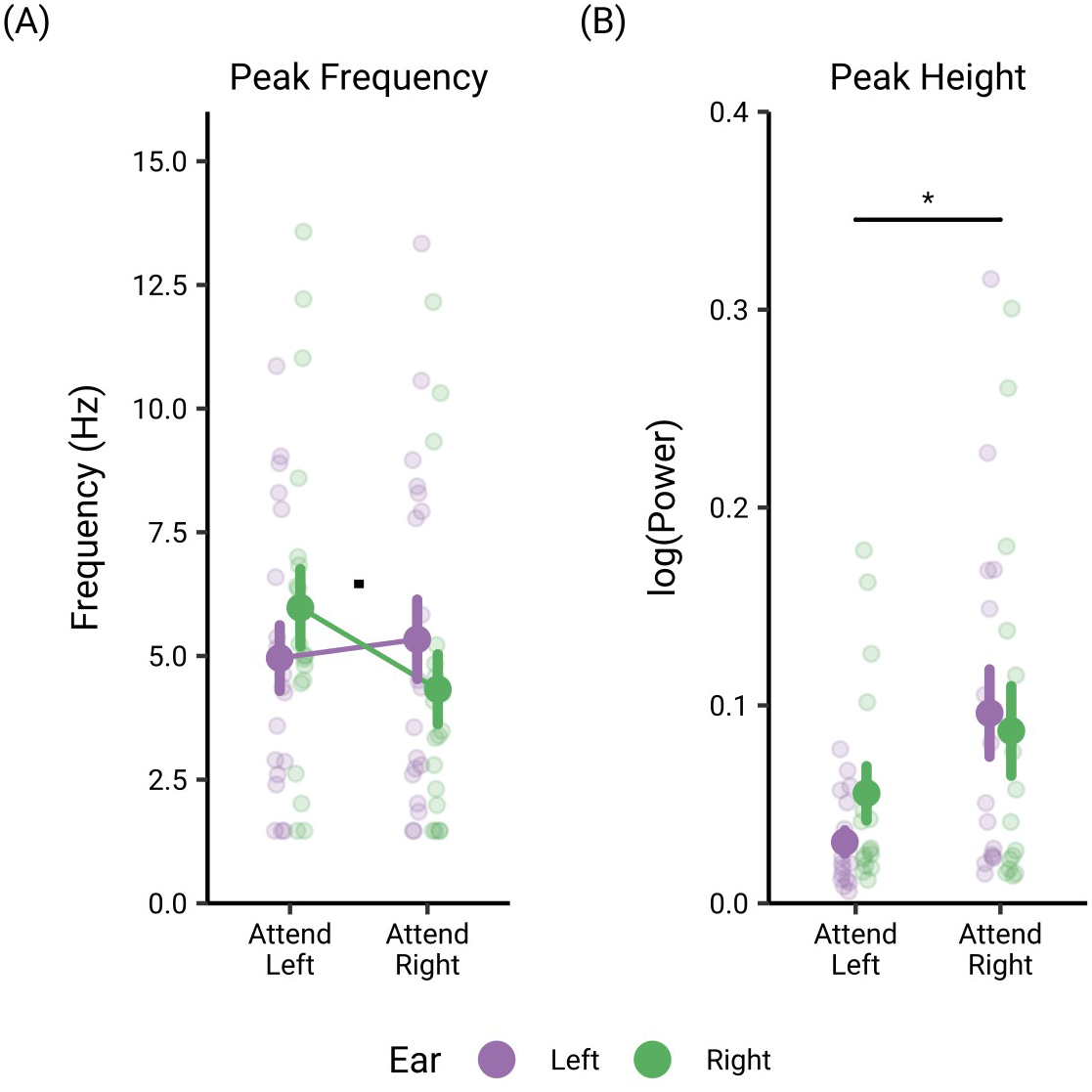
Peak analysis of OOA by FOOOF. (A) Results of a two factorial ANOVA (2×2) with the repeated measures factors ear (left & right) and attention condition (Attend Left & Attend Right) for peak frequency. On average theta rhythmicity for the OOA is revealed. The dot depicts the trend-level effect of the ordinal interaction. (B) Results of a two factorial ANOVA (2×2) with the repeated measures factors ear (left & right) and attention condition (Attend Left & Attend Right) for peak height of the peaks shown in (A).

The asterisk depicts the (on the 5%-level) significant main effect of attention condition. The big colored dots and error bars represent the mean and SEM.

Another aim of this study was to follow up on the relative increase of spectral power at frequencies of the cochlea’s theta rhythm when attention was shifted to the auditory modality as shown in Köhler et al. (2021). These findings depict a cochlear mechanism of selective attentional enhancement of the to-be-attended and deterioration of the to-be-ignored external stimulation for intermodal attention. Accordingly, we hypothesized in the context of interaural attention that the power at individual peak frequencies is increased in the to-be-attended compared to the to-be-ignored ear.

In addition to peak frequency, FOOOF reports the relative logarithmic power (peak height) above the aperiodic component for each identified peak. The average peak height in the left ear for the Attend Left condition was at *M* = 0.0309 (*SD* = 0.0237) and in the Attend Right condition at *M* = 0.0961 (*SD* = 0.0883). In the right ear the average peak height in the Attend Left condition was at *M* = 0.0556 (*SD* = 0.0547) and in the Attend Right condition at *M* = 0.0871 (*SD* = 0.0906). The result of a two factorial ANOVA (2×2) with the repeated measures factors ear (left & right) and attention condition (Attend Left & Attend Right) revealed no significant main effect for ear but, there was a significant main effect for attention condition (ear: *F*_(1, 15)_ = 0.29, *p* = 0.60; condition: *F*_(1, 15)_ = 6.95, *p* = 0.02). The interaction effect was not significant (*F*_(1, 15)_ = 1.34, *p* = 0.26). **Figure 2B** depicts the main effect of condition for peak height. The general increase in OOA peak power for the Attend Right condition in both ears could be explained by a general processing advantage for attention directed to the right side.

### Evoked Power at Stimulation Frequency Is Modulated by Interaural Attention

Subsequently, we tested if there are differences of the contralaterally evoked otoacoustic activity between attention conditions. For that, we separately calculated the evoked power of the envelope at the stimulation frequency during the omission periods for Attend Left and Attend Right trials. Two FDR-corrected dependent samples t-tests for each ear separately revealed a significant difference for the right ear (left ear: *T*_26_ = −1.07, *p* = 0.29; right ear: *T*_24_ = −2.80, *p* = 0.02). **Figure 3** illustrates the results of both t-tests. **Figure 3B** demonstrates that in the right ear the power for the Attend Left condition (*M* = 0.0113, *SD* = 0.0079) is higher than for the Attend Right condition (*M* = 0.0075, *SD* = 0.0050).

**Figure 3.**
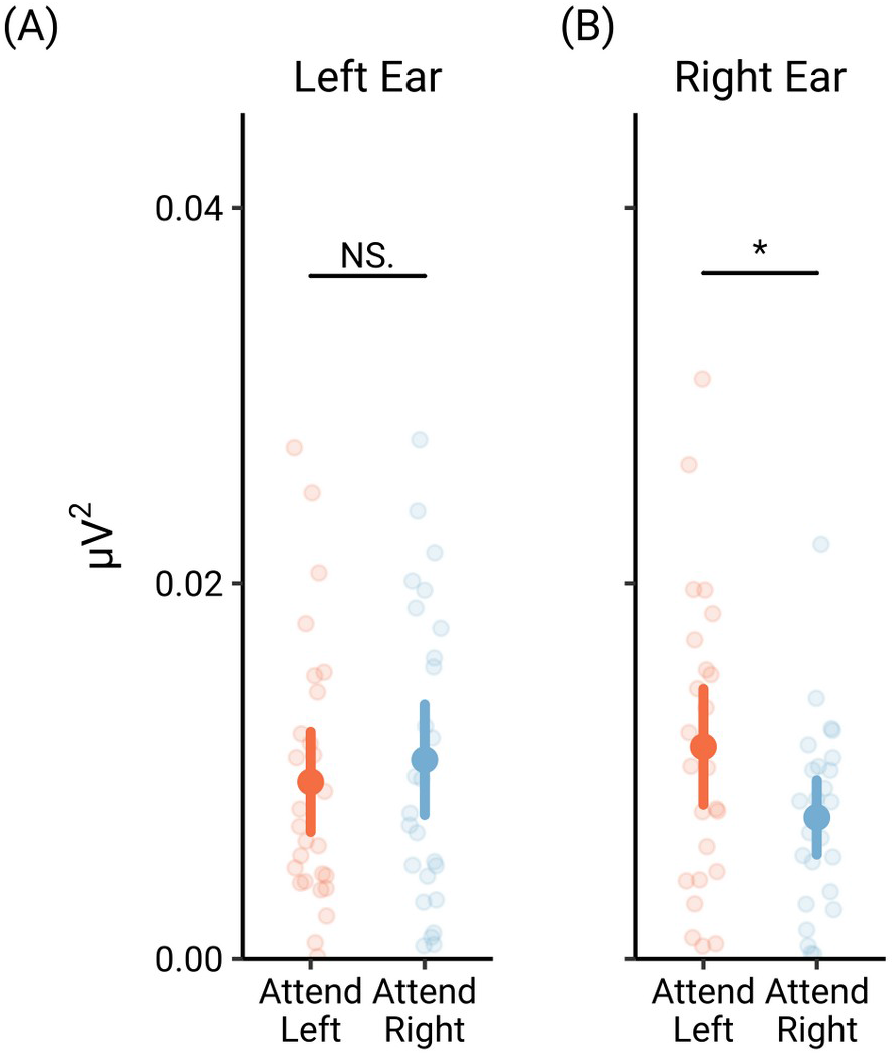
Evoked power at stimulation frequency (24 Hz) during omission periods. (A) Power of the two attention conditions for the left ear. (B) Power of the two attention conditions for the right ear.

**Figure 4.**
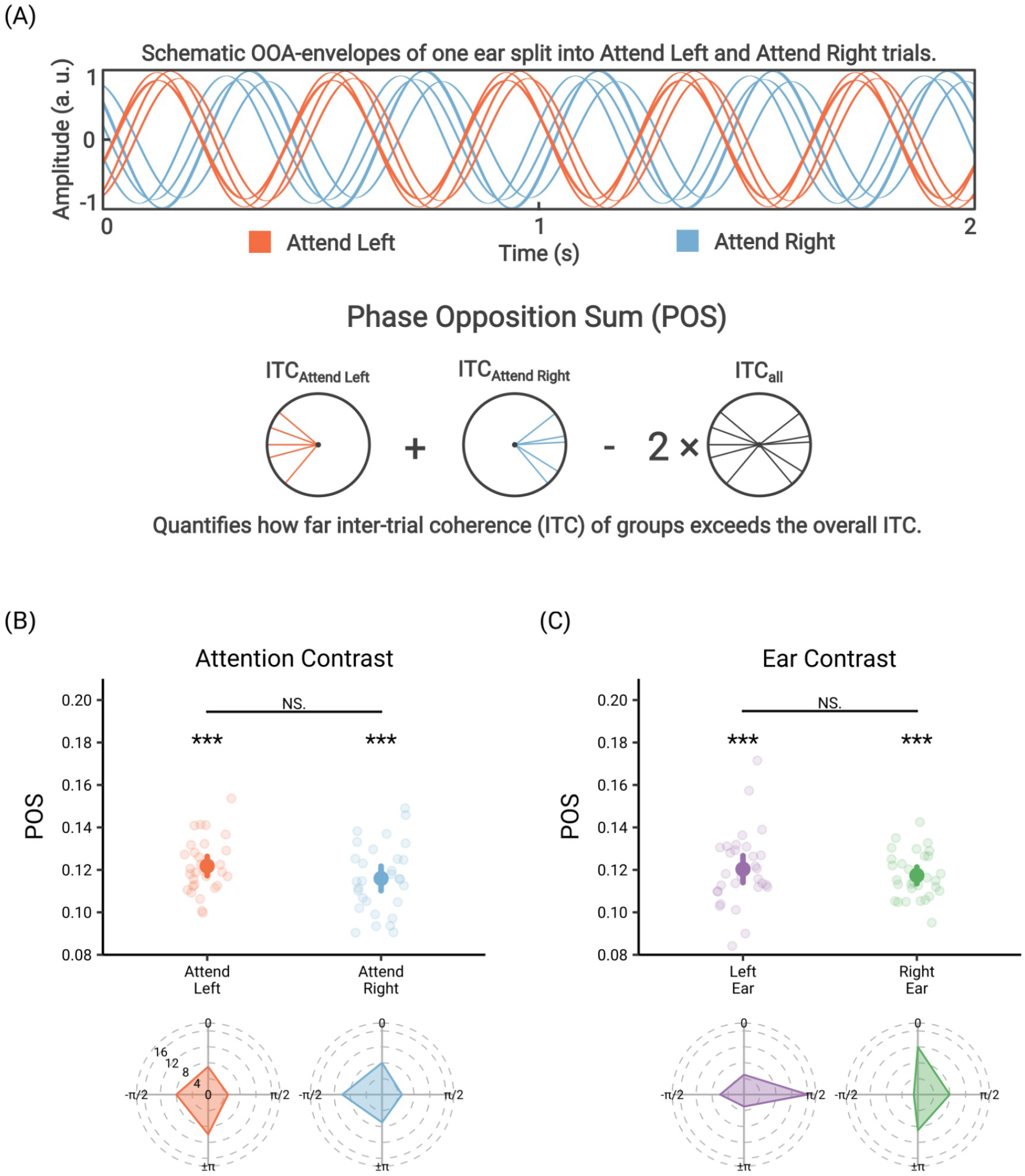
Rationale of phase analysis and cochlear phase effects during the silent cuetarget interval. (A) In the upper panel schematic OOA envelopes of one ear are illustrated. The envelopes are separately split into trials for the Attend Left and Attend Right condition. For illustration purposes the envelopes of both conditions oscillate almost at perfect antiphase. The consistency of the phase opposition between the Attend Left and Attend Right trials is calculated by the Phase Opposition Sum, whose formula is depicted in the lower panel. (B) POS for the ear contrasts. The POS between the left and right ear for all Attend Left trials is depicted in red and for all Attend Right trials in blue. The POS effects significantly differed from a null effect revealed by a permutation procedure. There was no difference between attention conditions. In the lower row the histograms for the relative phase differences are illustrated. The histograms are equally binned in four segments and frequencies are plotted at the lower bound of each bin. The phase of the left ear was normalized to 0. (C) POS for the attention contrasts. The POS between the Attend Left and Attend Right trials for the left ear is depicted in purple and for the right ear in green. The POS effects significantly differed from a null effect revealed by a permutation procedure. There was no difference between the ears. In the lower row the histograms for the relative phase differences are shown. The histograms are equally binned in four segments and frequencies are plotted at the lower bound of each bin. The phase of the Attend Left trials was normalized to 0.

All p-values are FDR-corrected. The asterisk depicts significance on the 5%-level. The big colored dots and error bars represent the mean and SEM.

### OOA Oscillates in Antiphase Between Ears and Attention Conditions

Besides attentional modulations in the power domain, we expected effects of interaural attention in the phase domain. In the cortical attention literature it is well established that the phases of neural oscillations affect how external stimuli are processed (e.g. stimuli arriving at the peak of a neural oscillation are in favor of further cortical processing than stimuli arriving at the trough). Attentional processes have the ability to alter or shift the phases of neural oscillations to favor some predicted to-be-attended stimuli over to-be-ignored stimuli. While power of cortical activity has been classically interpreted to indicate the level of excitation and inhibition, phase shifts reflect the timing regulation of excitability. Here, we aimed to investigate if phase shifts of the OOA represent an analogous mechanism already at the level of the cochlea. We hypothesized that the phase during the cue-target interval in the same attention condition is different between the two ears (ear contrasts). Moreover, we speculated that the phase in the same ear is different between both attention conditions (attention contrasts).

The big dots and error bars represent the mean and SEM. The three asterisks illustrate significance on the 0.001%-significance level.

We analyzed phase differences by calculating the POS for every contrast of interest. The POS is a measure to calculate the consistency of phase opposition between two signals. **Figure 4A** illustrates the rationale of this analysis.

For testing the first hypothesis (ear contrasts) we selected all Attend Left trials and contrasted the signal from the left with the right ear. After that we did the same for all Attend Right trials. Altogether, we compared the phases of the to-be-attended ear with the to-be-ignored ear in both conditions. As we were interested in phase differences at frequencies of the cochlear rhythm, we averaged the POS values between 1.56 and 9.82 Hz. The limits for this frequency range were defined by ± 1 *SD* of the mean peak frequency from all peaks that were previously identified. While attention was shifted to the left side the POS between the two ears was *M* = 0.122 (*SD* = 0.013, *p* = 5.76e-13) and while it was shifted to the right side it was *M* = 0.116 (*SD* = 0.016, *p* = 1.1e-08). There was no significant difference between both contrasts (*T*_(29)_ = 1.57, *p* = 0.13). **Figure 4B** shows the reported effects of the POS for the ear contrasts. The two panels in the lower row illustrate the histograms of the relative phase differences for the Attend Left and Attend Right trials while the phase of the left ear was normalized to 0. The histograms are equally binned in four segments and frequencies are plotted at the lower bound of each bin.

Next, we tested the second hypothesis (attention contrasts) by applying the same approach as for the first one with only one difference. We contrasted for the left ear the signal from Attend Left trials with the signal from Attend Right trials. After that we did the same for the right ear. We then compared the phases of one ear while it was to-be-attended and while it was to-be-ignored. In the left ear the overall POS between the both attention conditions was *M* = 0.120 (*SD* = 0.018, *p* = 9.19e-10) and *M* = 0.117 (*SD* = 0.011, *p* = 3.09e-11) in the right ear. There was no significant difference between both contrasts (*T*_(29)_ = 0.76, *p* = 0.45). **Figure 4C** visualizes the effects of the POS for the attention contrasts.

In order to provide additional evidence that phases are modulated by interaural attention we employed the same analysis approach to the phases from the omission periods of the click trains. In this way, we were able to assess phase differences of the cochlear rhythm during the presentation of the target. The POS for the ear contrasts was *M* = 0.223 (*SD* = 0.155, *p* = 0.006) for Attend Left trials and *M* = 0.222 (*SD* = 0.159, *p* = 0.010) for Attend Right trials (See **Figure 5A**). The POS for the attention condition contrasts was *M* = 0.203 (*SD* = 0.185, *p* = 0.026) in the left ear and *M* = 0.242 (*SD* = 0.192, *p* = 0.011) in the right ear (See **Figure 5B**). This analysis indicates that there is phase opposition during task execution for both the attention condition and the ear contrasts. Hereby, these results corroborate that the phase opposition effects during the cue-target interval were induced by top-down attentional processes.

**Figure 5.**
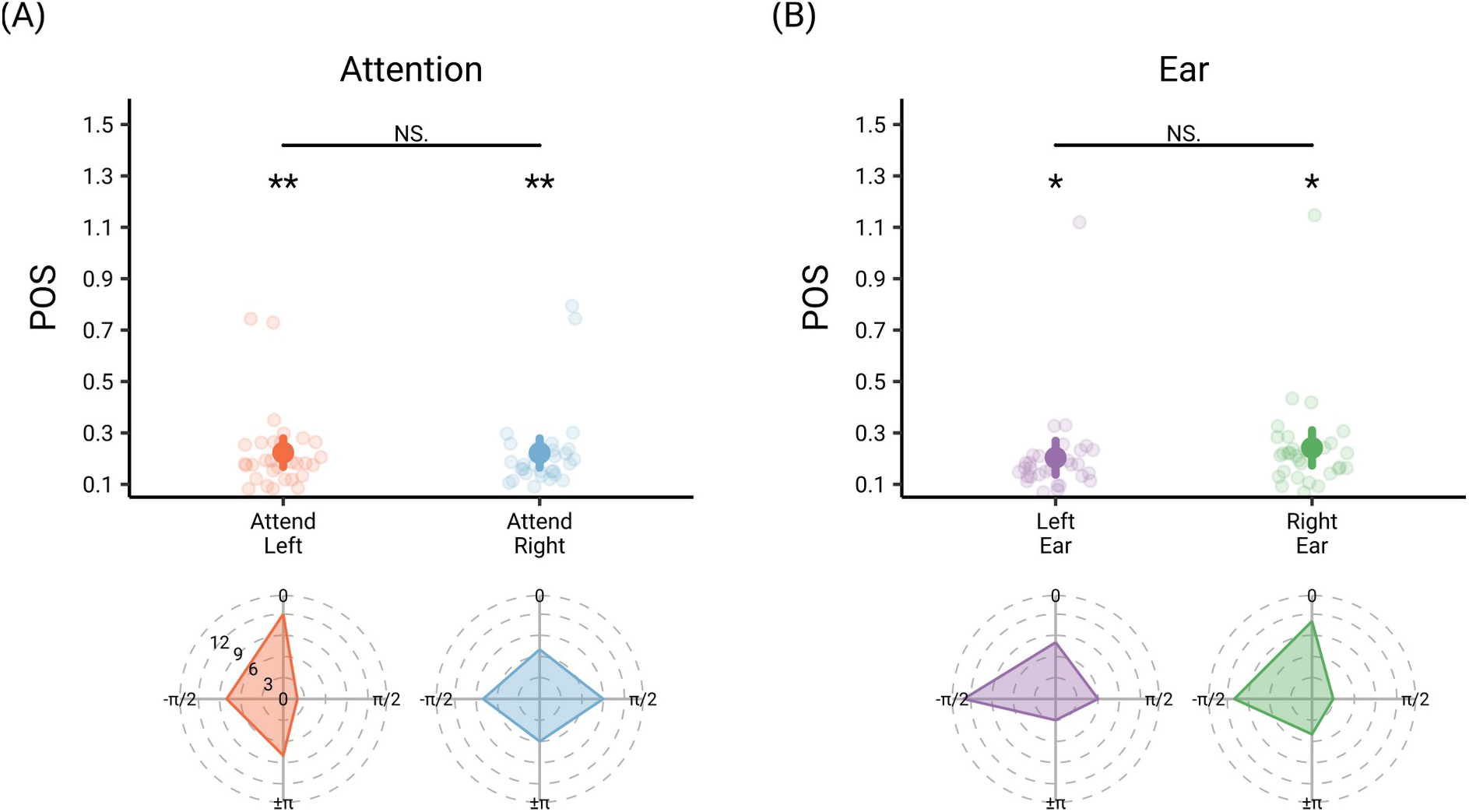
Cochlear phase effects during the omission periods. (A) POS for the ear contrasts. The POS between the left and right ear for all Attend Left trials is depicted in red and for all Attend Right trials in blue. The POS effects significantly differed from a null effect revealed by a permutation procedure. There was no difference between attention conditions. In the lower row the histograms for the relative phase differences are illustrated. The histograms are equally binned in four segments and frequencies are plotted at the lower bound of each bin. The phase of the left ear was normalized to 0. (B) POS for the attention contrasts. The POS between the Attend Left and Attend Right trials for the left ear is depicted in purple and for the right ear in green. The POS effects significantly differed from a null effect revealed by a permutation procedure. There was no difference between the ears. In the lower row the histograms for the relative phase differences are shown. The histograms are equally binned in four segments and frequencies are plotted at the lower bound of each bin. The phase of the Attend Left trials was normalized to 0.

The big dots and error bars represent the mean and SEM. One asterisk illustrates significance on the 0.05%-significance level and two asterisks on the 0.01%-level.

All in all, the reported results show that the phase of the oscillatory activity at the level of the auditory periphery is modulated by attention processes. Importantly, the results of the attention contrasts provide conclusive evidence that supports this notion. While phase opposition between both ears could be an adaptation to (cochlear) physiological processes, phase opposition in one ear between both attention conditions allows one to draw the conclusion that attention modulates the phase of OOA.

### OOA Is Not Phase Locked to Eye Movements

Gruters et al. (2018) demonstrated that the eardrums oscillate in relation to horizontal eye movements. These so-called eye movement-related eardrum oscillations (EMREOs) occur predominantly between 20 - 40 Hz of the acoustic spectrum measured from the ear canals. Our reported effects of phase opposition are at much higher acoustic frequencies (1500 - 2000 Hz). However, via phase-amplitude coupling the envelopes at high frequencies could still be confounded by low frequency eye movement-related acoustic activity during the cue-target interval. If this was the case, then a clear phase coupling between OOA and eye movements would be predicted. In order to test this possibility we calculated the PLV between the eye movements (ocular components from EEG data) and the acoustic envelope from both ears at the same frequencies as was used for the POS analysis.

In three subjects no eye movement-related ICA component could be identified as these subjects did not show marked eye movements in the EEG-data during the cue-target period. The PLV in the left ear when attention was directed to the left side was *M* = 0.058 (*SD* = 0.004, *p* = 0.289) and in the right ear *M* = 0.057 (*SD* = 0.004, *p* = 0.852). The PLV in the left ear when attention was directed to the right side was *M* = 0.061 (*SD* = 0.006, *p* = 0.197) and in the right ear *M* = 0.057 (SD = 0.007, *p* = 0.573).

We failed to show significant phase locking to eye movements in the left and right ear for both attention conditions. Thus, the ocular process that underlies the previously reported EMREOs (Gruters et al., 2018) is only a very weak candidate explanation for our attentional OOA effects.

## Discussion

Previous research on top-down modulations of the phase of auditory perception was focused on cortical and behavioral effects but neglected the cochlea, which is the most peripheral part of the auditory system and innervated by efferent connections arising from the primary auditory cortex. In our previous study we reported a putatively endogenous theta-rhythmic pattern of otoacoustic activity (Köhler et al., 2021). While low frequency power of oscillatory cochlear activity was modulated by topdown intermodal attention its frequency and phase was independent of it. As the paradigm of the previous study was not suitable for assessing top-down modulations of phase, we implemented an interaural attention paradigm that allowed us to draw conclusions on how the phases of the cochlea’s theta rhythm in both ears are modulated by attention processes. We show that interaural attention consistently modulates the phases in both ears. In the past, such effects were only reported for cortical signals and behavioral performance. To the best of our knowledge this is the first time that phase modulations of the auditory receptor are reported.

### Cochlear Acoustic Activity Is Theta-rhythmically Modulated

Our analysis of the OOA during the cue-target interval replicated the findings from our previous study, namely the existence of a theta-rhythmic (~5 Hz on average) modulation of the cochlea’s otoacoustic activity (Köhler et al., 2021). Thereby, aperiodic (“1/f”) contributions to this effect were ruled out. This finding is in line with the results from a study that investigated oscillations of behavioral performance in a bilateral pitch-identification task (Ho et al., 2017). The authors reported an oscillation of behavioral performance in the theta (~6 Hz) and low alpha (~8 Hz) range. In addition, a second study that adapted the experimental paradigm from Ho et al. (Ho et al., 2017) could also report a theta-rhythmic modulation of auditory behavioral performance (Plöchl et al., 2021). Considering the anatomical structure of the efferent auditory system (Terreros & Delano, 2015), cochlear rhythmicities could be modulated by cortical (attention) processes or be adaptations to physiological processes of the auditory receptor.

### Cochlear Acoustic Activity Oscillates in Antiphase Between Ears and Direction of Attention Within an Ear

The temporal pattern of the cochlear rhythmic activity during the cue-target interval displayed significant phase opposition at frequencies encompassing the theta-band. This phenomenon was not only present between the to-be-attended and to-be-ignored ear but also in the same ear between both attention conditions. These results are very similar to that observed in the visual and recently emerging auditory rhythmic sampling literature for both neural activity and behavioral performance, expanding it to the level of the auditory receptor (Fiebelkorn et al., 2013; Ho et al., 2017; Kayser, 2019; Kubetschek & Kayser, 2021; Landau & Fries, 2012; Plöchl et al., 2021; Spyropoulos et al., 2018). Moreover, on the basis of the PLV analysis it seems very unlikely that these effects are driven by eye movement-related activity.

For visual attention it is established that simultaneously presented objects are sampled in sequence at ~4 Hz (Fiebelkorn et al., 2013, 2018; Helfrich et al., 2018; Landau et al., 2015; Landau & Fries, 2012). For the auditory modality studies reported conflicting results of such effects (Ho et al., 2017; İlhan & VanRullen, 2012; Ng et al., 2012; Plöchl et al., 2021; Zoefel & Heil, 2013). Either effects were absent (İlhan & VanRullen, 2012), at least partly dependent on the power and phase of ongoing neural theta oscillations (Ng et al., 2012), or attributed to artifacts (Zoefel & Heil, 2013). Regardless, recent studies were consistently able to demonstrate rhythmic modulations of behavioral detection performance (Ho et al., 2017, 2019; Kayser, 2019; Plöchl et al., 2021). For instance, Ho et al. (2017), who applied signal detection theory to test for oscillations of behavioral performance in a bilateral pitch-identification task, demonstrated that both criterion and sensitivity oscillate in antiphase between the left and right ear. These findings are in line with the phase opposition of OOA between the to-be-attended and to-be-ignored ear reported here. However, such an effect does not rule out the possibility that both ears endogenously sample auditory input in antiphase. Certainly, OOA also demonstrates phase opposition between both attention conditions in the same ear. This phenomenon provides evidence that the auditory streams from both ears are not endogenously sampled in antiphase but interaural attention systematically modulates the temporal dynamics of auditory sampling. Together, both effects corroborate the existence of alternating attentional states directly affecting cochlear processes.

### Interaural Attention Modulates the Timing of Excitation & Inhibition of Cochlear Activity

In Köhler et al. (2021) we reported the existence of a systematic variation of OOA level but not phase for intermodal attention. Interestingly, in the current study interaural attention systematically modulates the phase of OOA but against our hypothesis not its level. Thus, it seems that both types of attention impact cochlear activity in a differential manner. While intermodal attention modulates the level of cochlear activity, interaural attention modulates the timing. In situations where the former is of relevance the overall auditory input is either distracting and should be ignored or relevant and processing should be facilitated. A mechanism affecting the level of amplification of auditory input at the level of the cochlea seems to fit this aim. Conversely, in situations where interaural attention is of relevance we initially hypothesized that it deploys additionally to cochlear level differences between ears also differences in the temporal organization of the auditory input from both ears. Our current results revealed that interaural attention relies only on a mechanism affecting the timing of cochlear activity. In this regard, on the cochlear level intermodal attention seems to manifest via level differences of cochlear activity and interaural attention via its temporal orchestration.

So far, there is only one study that systematically investigated effects of interaural attention on the acoustic activity of the outer hair cells. Srinivasan et al. (2014) recorded distortion product otoacoustic emissions (DPOAE) while participants had to identify brief tones in the ipsi- or contralateral ear or brief phase shifts of a visual grating. Interestingly, DPOAE levels were smallest when attention was shifted to the ipsilateral ear, where DPOAEs were recorded from, and highest for visual attention.

These results stand in contrast to widely accepted effects observed by electrophysiological measures of peripheral auditory function (Delano et al., 2007; Lukas, 1980), otoacoustic (Dragicevic et al., 2019; Köhler et al., 2021; Wittekindt et al., 2014), and cortical measures (Johnson & Zatorre, 2005; Kauramäki et al., 2007; Woldorff et al., 1987). However, the authors explain their found effect by the fact that participants were instructed to focus the DPOAE primary tones, which are some cochlear distance apart, and by the tonotopic tuning of the MOC the response of the DPOAE could be suppressed.

In the current study the OOA did not display a systematic variation in peak height but phase between ears and attention conditions. In line with our results there is evidence from studies on dichotic listening while recording physiological noise from the ear canal that there is no difference in noise level between the ipsi- and contralateral ear (Walsh et al., 2014, 2015). These results speak for interaural attention manifesting via the timing aspect of cochlear activity. In contrast to the current study, the paradigm of Srinivasan et al. (2014) did not exclusively assess interaural attention but also intermodal attention. That is, participants shifting their attention to one ear consistently had to not only ignore auditory input from the other ear but also visual input. However, their results support our proposition that intermodal attention manifests via level differences of otoacoustic activity.

### The Input from Both Ears Is Perceived as Two Independent Objects Rather than One Object with Two Locations

Lately one study suggests that visual and auditory oscillations share a general attentional mechanism, which theta-rhythmically samples target locations and objects, respectively (Plöchl et al., 2021). Thereby, supramodal attentional sampling switches between two objects at a rate of ~ 4 Hz and between two target locations within a single object at ~8 Hz. Interestingly, Plöchl et al. (2021) found a significant phase opposition only at ~8 Hz for auditory detection performance. The authors attributed that to the fact that auditory input from both ears may not consistently be perceived as two independent objects but rather as a single object containing two target locations. Also the from Ho et al. (2017) reported effects of behavioral oscillations pointing more towards ~8 Hz than ~4 Hz seem to support this assumption. In contrast, our results of cochlear oscillations at ~5 Hz speak more for the perception of two auditory streams as two independent objects. However, the question to what extent such oscillatory effects of behavior are represented on a cochlear level is a matter for future research.

### Aspects for Future Studies

The rhythmic modulation in our previous study was not affected in frequency by intermodal (auditory/visual) attentional processes. Hence, we argued for a general endogenous cochlear rhythm. However, the trend-level effect of frequency reported here could point to a systematic modulation of the frequency of the cochlear rhythmic activity by interaural attention processes. More precisely, a reduced frequency for the to-be-attended compared to the to-be-ignored ear. This phenomenon could be interpreted in line with the active sampling literature proposing two theta-dependent states of attention that either facilitate sampling of an object/location or the likelihood of a shift to another object/location (Fiebelkorn & Kastner, 2019). In the current study there were two locations as auditory stimulation was fully lateralized to the left and right ear. Therefore, for active listeners occasional shifting to the to-be-ignored ear would have likely happened. An increase in frequency for the to-be-ignored ear would lead to a reduction in time required to occasionally sample this ear. Again, it has to be noted that this effect did just fail to reach statistical significance. Nevertheless, it would be interesting for future studies to address the existence of this phenomenon.

## Conclusion

Here we demonstrate that the phase of cochlear theta-rhythmic activity is systematically modulated by interaural attention. In doing so, the present results suggest that cueing events can orchestrate endogenous cochlear oscillations to putatively affect processing of upcoming stimuli. In addition, this study not only adds to a growing body of literature providing evidence that attentional sampling is not restricted to the visual modality but also extends this mechanism to the most peripheral stage of the auditory efferent system. In the context of the corticofugal pathways we provide an additional link between the primary auditory cortex and behavior. Yet, at this point it remains an open question if (1) perceptual and attentional rhythmicities are genuinely driven by cortical effects or are an adaptation to physiological properties of the cochlea, (2) oscillatory effects of behavior are reflected on a cochlear level. Future studies should investigate the relationship between cochlear, (auditory) cortical, and behavioral oscillations. Moreover, it would be interesting to study the mechanistic properties of this rhythm in hearing impaired individuals.

## Acknowledgments

We thank Konstantin Kölle for his help with the measurements and Daniel Walker for proofreading the manuscript. Moreover, we are grateful to Dr. Gianpaolo Demarchi for discussions and assistance concerning this project. This work was supported by the DOC Fellowship Programme of the Austrian Academy of Sciences.

